# Optogenetic mapping of rhythmic phase-dependent excitability in the mouse striatum

**DOI:** 10.1101/2024.02.01.578473

**Authors:** Manish Mohapatra, J. Eric Carmichael, Kyle S. Smith, Matthijs A. A. van der Meer

## Abstract

The striatum is thought to switch flexibly between multiple converging inputs to support adaptive behavior. The “communication through coherence” (CTC) hypothesis is a potential mechanism to implement such a flexible switching. For CTC to work in the striatum, striatal excitability must show rhythmic fluctuations, such as those related to the phase of the striatal local field potential (LFP). To test this fundamental requirement, we delivered a constant input stimulus to ChR2-expressing striatal fast-spiking PV+ interneurons (FSIs) in head-fixed awake mice (PV-Cre:Ai-32, n = 18, 9 female), and determined whether the response to this stimulus varied with LFP phase. We found that approximately one-third (41.2%) of FSIs exhibited significant phase-dependent excitability in at least one LFP frequency band. Phase-dependent excitability was most prominent in the delta (2–5 Hz) frequency band, both in terms of prevalence (23.5% of FSIs sampled) and magnitude (phase modulation strength: 22% of average response). The most excitable phase tended to align with endogenous phase-locking, again most clearly in the delta band. These results bolster the functional relevance of the striatal field potential and spike-field relationships, and provide proof-of-principle support for the possibility of CTC in the striatum.

**Significance Statement:** The striatum is thought to switch dynamically between multiple converging inputs. A leading idea for how this is accomplished is through communication through coherence (CTC). A fundamental, but previously untested, requirement for CTC to work is that striatum must show changes in excitability that depend on local field potential phase. We find that about one-third of striatal neurons show phase-dependent excitability, providing proof of principle for CTC-like switching in striatum.

## Introduction

The striatum, and its ventral aspect in particular, receives converging inputs from a number of different brain areas, such as frontal cortex, hippocampus, thalamus and the amygdala, which it is thought to flexibly select between in order to support adaptive behavior. For instance, in some tasks, contextual information from the frontal cortex or hippocampus disambiguates whether a given cue has motivational relevance or not (Humphries & Prescott, 2010; Floresco, 2015). At the cellular level, such “gating” or “switching” is thought to be implemented in so-called “up-” and “down-states” which can be measured intracellularly (O’Donnell & Grace, 1995). However, it is unclear to what extent this single-cell gating mechanism shown in anesthetized preparations generalizes to interactions between populations of neurons that span different brain structures in awake animals.

A leading proposal for how flexible gating between different inputs is implemented in populations of neurons is “communication through coherence” and related ideas (CTC; Fries, 2005, 2015; Lakatos et al., 2008; T. Akam & Kullmann, 2010; Busch et al., 2009; Buzsáki & Schomburg, 2015). CTC postulates that a downstream “receiver” population exhibits fluctuations in excitability (that is, the likelihood of generating a response for an input of a given size) such that inputs that arrive at the time of maximum excitability are getting their message transmitted to the receiver, whereas inputs that arrive at times of low excitability are getting their message blocked. Thus, the most effective inputs consistently arrive at times of high excitability; those inputs are said to be *coherent* with the receiving area. Substantial support for CTC has come from the demonstration that coherence, as measured by local field potentials (LFPs), relates to information transfer, most prominently in the visual system (Bosman et al., 2012) and hippocampus (Colgin et al., 2009).

A number of studies have documented the existence of widespread oscillations across single cell and field potential levels not only in striatum itself but also in essentially every major input to the striatum (Leung & Yim, 1993; Berke et al., 2004; Brown & Williams, 2005; Howe et al., 2011; Von Nicolai et al., 2014; Feingold et al., 2015; van der Meer, 2010). Moreover, these oscillations dynamically synchronize depending on frequency, task, and brain state (DeCoteau et al., 2007; Popescu et al., 2009; Gruber et al., 2009; Hunt et al., 2009; van der Meer & Redish, 2011; Cohen et al., 2012; Lemaire et al., 2012; Dejean et al., 2013; Lansink et al., 2016; Catanese et al., 2016; Amadei et al., 2017) in a manner highly suggestive of CTC. However, a foundational and as yet untested requirement of striatal CTC is that the receiving area must show fluctuations in excitability. If a downstream target’s excitability does not vary, then coherence between the input and the target does not allow for a way for the input timing to drive the target any more effectively compared to when the input and target phases are not coherent. In other words, communication can occur through coherence *only if* the downstream region shows fluctuations in excitability. Whether this basic requirement for CTC is met in the striatum is unknown.

In this study, we test the hypothesis that striatal excitability (i.e. the response to a constant input) varies with the phase of oscillations in the striatal field potential^1^. Fluctuations in excitability are hard to measure directly, especially *in vivo*, because they are realized in intracellular quantities such as the subthreshold membrane potential and the state of various ion channels such as those responsible for shunting inhibition (Mitchell & Silver, 2003). To get around these challenges, we apply optical stimulation to channelrhodopsin-expressing striatal neurons at various phases of the ongoing striatal LFP, and test whether the response magnitude depends on the LFP phase at the time of stimulation. Phase-dependent excitability will manifest as systematically higher responses when the optical stimulation aligns with the preferred LFP phase: this result would constitute key support for CTC. Conversely, if the response magnitude is unaffected by the LFP phase at the time of the stim, there are no fluctuations in excitability, and CTC cannot occur (see **Figure 1** for a schematic illustration of this idea). Briefly, we find that ∼41% of striatal fast-spiking interneurons exhibit phase-dependent excitability, constituting proof-of-principle for a fundamental requirement of CTC.

**Figure 1:**
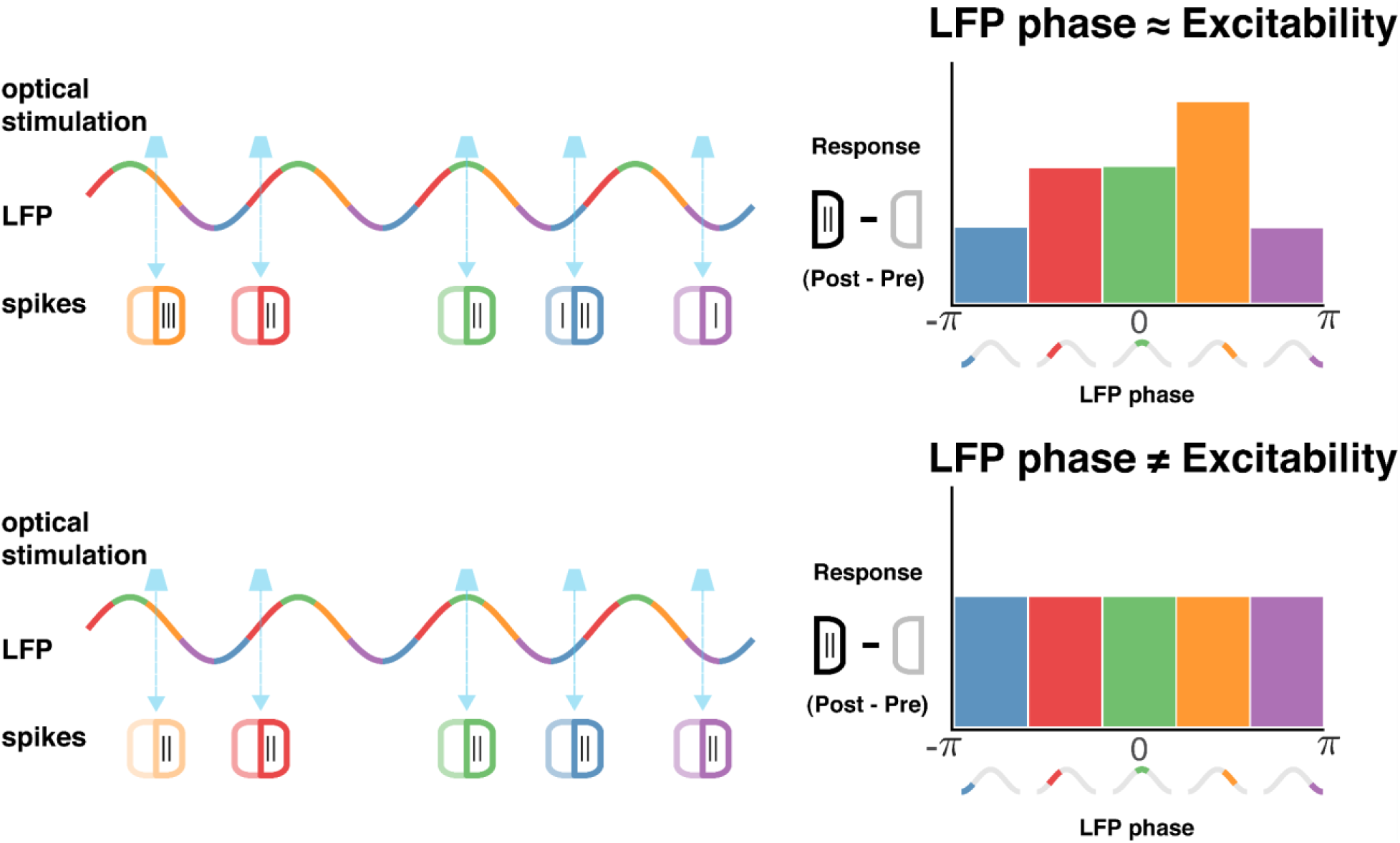
Schematic of predictions following from the hypothesis that striatal local field potential oscillations reflect local neuronal excitability (top) and the null hypothesis that excitability is unrelated to the field potential (bottom). Local field potential (LFP) oscillations are depicted as sinusoidal traces with their phases starting from the valley (-π radians; negative LFP voltage), rising to the peak (0 radians; positive LFP voltage) and finally ending at the valley again (π radians), divided into 5 color-coded bins (left). Randomly timed optical stimulation of constant intensity (cyan arrows) occurs across different phases, evoking spikes in channelrhodopsin-expressing neurons. The evoked response is quantified as the difference between the number of post-stimulus and pre-stimulus spikes (“post minus pre”). If a neuron’s excitability is related to the LFP oscillation phase, then its response magnitude (post-stimulus minus pre-stimulus spike count) to optical stimulation should be highest when stimulation aligns with that neuron’s most excitable phase (green in this example; top right). In contrast, if LFP phase is unrelated to excitability, then response magnitude should not change as a function of stimulus LFP phase (bottom right).

We also test the secondary hypothesis that the most excitable LFP phase aligns with the phase to which neurons preferentially phase-lock endogenously. In general, LFPs reflect summed transmembrane currents, such that events highly synchronous in time (e.g. a burst of feedforward input) and in space (e.g. that input activates many synapses) result in the largest LFP deflections (Buzsáki et al., 2012). Although the specific LFP phase that corresponds to membrane depolarization and increased spiking often depends critically on recording location (e.g. the phase of hippocampal theta reverses 180° across the cell layer), LFP phase carries information about spiking, as demonstrated by the widespread nature of endogenous spike-field relationships (e.g. phase locking, including in the striatum: Sharott et al., 2009 for dorsal striatum; van der Meer & Redish, 2009; Berke, 2011 for ventral striatum). From the perspective of CTC, what is required is not just spike-field locking, but a change in how effective a given input is in causing downstream spiking (i.e. excitability). The most effective phase for CTC may, or may not, align with the preferred phase-locking phase: if a neuron is already driven hard by endogenous inputs, then additional input may be ineffective due to low input resistance, adaptation, and/or refractoriness (Cannon and Miller, 2016). Alternatively, if a neuron is strongly inhibited (e.g. in a down-state) then inputs may also be ineffective. Thus, it is an empirical question what the preferred LFP phase for excitability is, and how it relates to the preferred LFP phase for endogenous phase-locking. Determining this relationship is a prerequisite for knowing whether the commonly observed phenomenon of LFP coherence (synchrony) between striatum and its inputs can be interpreted as conducive, or prohibitive, for communication.

## Materials and Methods

### Subjects and timeline

PV-Cre female mice were bred with Ai-32 (ChR2(H134R)-EYFP) males (Jackson Labs), to produce offspring PV-Cre:Ai-32 mice expressing ChR2 in parvalbumin-positive (PV+) interneurons throughout the brain, including the striatum (Figure 2A). These PV-Cre:Ai-32 mice (n = 18, 9 female) were aged between 2 and 14 months and were housed individually for at least 1 week prior to surgery. All surgeries were performed under general anesthesia (isoflurane) and analgesia (ketamine) in accordance with Dartmouth IACUC protocol 00002235; see Gmaz & van der Meer (2022) for further details. Briefly, custom titanium head-bars were attached to the skull using dental cement (Parkell Metabond) and craniotomy locations over the striatum (AP: bregma +0.8 ‒ 1.4 mm; ML: bregma ±1.0 ‒ 2.0 mm) marked. Following a recovery period of 2 days, mice were habituated over 3‒6 days to spend up to 1 hour in the experimental setup where they were free to run on a running wheel while being head-fixed. A break of 2 days was given after this period. In a second surgery, bilateral craniotomies were then drilled over the striatum, and 2 stainless steel screws were placed in the cortex bilaterally (AP: bregma −1.5mm, ML: bregma ±1 mm) to serve as ground and reference electrodes. Another recovery period of 2 days was provided before recording sessions (2‒11 per animal, 1 session per day) began. Following the final recording session, animals were euthanized (isoflurane overdose), perfused, and their brains recovered for histological processing.

**Figure 2:**
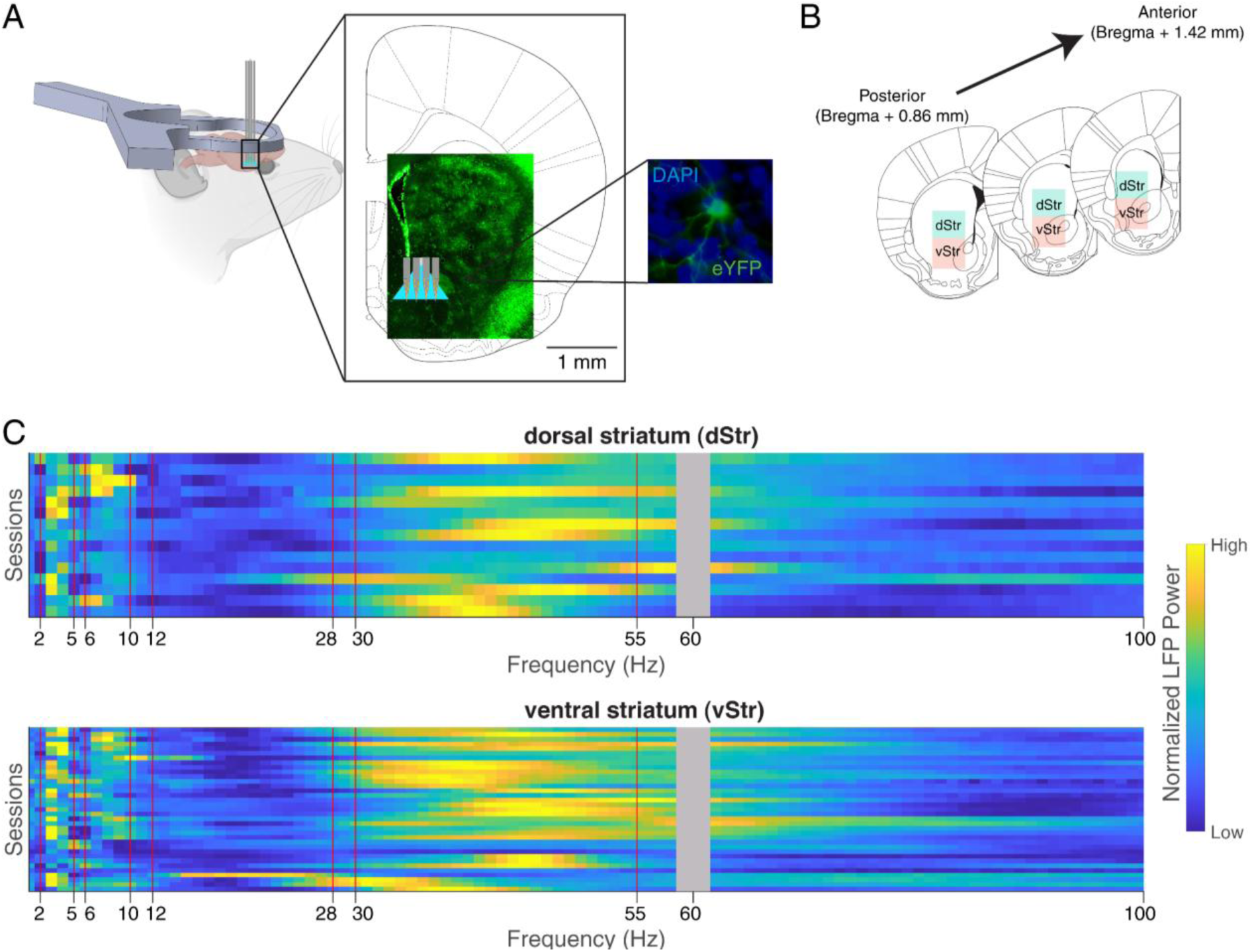
Schematic of experimental setup and recording locations. **A**: PV-Cre x Ai-32 mice (n = 18, 9 female) received surgically affixed headbars on their skulls and craniotomies over the striatum. These mice expressed ChR2 (EYFP) in PV+ neurons across the brain, including the striatum, as visualized in a representative brain slice under fluorescence microscopy (inset). During each recording session, we lowered an optrode into a craniotomy targeting either the dorsal striatum (dStr) or the ventral striatum (vStr) for simultaneous electrophysiological recording and optical stimulation with blue light. **B**: Recording locations were determined based on a combination of craniotomy coordinates, recording depths, and histology, and classified as either from dorsal striatum (dStr, green square, 2.5‒3.5 mm deep), or ventral striatum (vStr, red square, 3.5‒4.5 mm deep). **C:** Power spectral densities (PSDs) of the LFP channels in all included sessions, i.e. those that had opto-responsive cells, displayed as heatmaps for dStr **(top)** and vStr **(bottom)**. Each row denotes a single session’s PSD which has been min-max normalized after subtracting away its fractal component obtained via IRASA (Oostenveld et al., 2011; Wen & Liu, 2016). Thus, lighter colors denote peaks in the PSD which lie above the broadband (1/f) activity, indicating the presence of oscillations at those frequencies. Consistent peaks across sessions were seen in the 2‒5 Hz, 6‒10 Hz, and 30‒55 Hz range (bounded by red vertical lines), thus these were selected as the frequency bands of interest, along with the beta band (12–28 Hz) due to its theoretical importance.

### Experimental design

The overall aim of this study was to examine the relationship between the excitability of striatal neurons and the phase of ongoing striatal LFP oscillations. To this end, we used optical stimulation in mice expressing ChR2 in PV+ interneurons. By using a large number of stimulations (1500) with a variable time interval between them (0.5‒3.5 sec), we uniformly sampled the phase range of oscillations in the major striatal LFP frequency bands (delta: 2‒5 Hz, theta: 6‒10 Hz, beta: 12– 28 Hz, gamma: 30‒55 Hz). We targeted PV+ interneurons because these neurons exhibit stronger oscillatory properties compared to other cell types (van der Meer & Redish, 2009; Kalenscher et al., 2010; Sharott et al., 2012; van der Meer et al., 2019) and exert widespread and powerful influence over striatal projection neurons (“gain control”; Gittis et al. 2010; Owen et al. 2018). As a practical benefit, PV+ neurons are relatively scarce among the striatal neuronal population (1‒ 5%, less in the ventral compared to dorsal striatum; Tepper & Bolam, 2004; Tepper et al., 2010), minimizing the possibility of stimulation triggering large-scale network effects that could interfere with spike-sorting and LFP phase estimation.

### Experimental setup and recording procedure

Acute recordings were performed using multi-shank silicon probes equipped with an optic fiber whose tip was positioned near the recording sites on the middle shanks. The specific probe configuration and light source varied depending on probe and equipment availability (Neuronexus Buz32 probe with 50μm 0.39 NA optic fiber, driven by a Shanghai Dream 473nm blue laser, Uniblitz LS2 Optomechanical shutter and Thorlabs 125 um/0.39 NA optical patch cord; alternatively, a Cambridge Neurotech Assy 37-P or 77-P1 probe with 200 um 0.5 NA optic fiber, Prizmatix 460nm Blue LED driver and Thorlabs 500um/0.5 NA optical patch cord) but these different configurations produced equivalent data and were pooled for all analyses. A Master-8 (AMPI) or PulsePal v2 (Sanworks) controlled both the timing and width of the light pulses used for optical stimulation.

Silicon probe signals were preamplified (HS-36, Neuralynx) and acquired using a Digital Lynx acquisition system (Neuralynx). Before each day’s recording, the power output at the tip of the optic fiber was measured using a Thorlabs PM1000 USB power meter and a Thorlabs S140C sensor. Light output was calibrated to an analog output dial on the light source, enabling power level control between 1‒10 mW in 1 mW steps. Following this, the animal was secured in a custom built head-fixed setup atop a running wheel (Gmaz & van der Meer, 2022).

Following calibration, the optrode was positioned into a selected craniotomy and then lowered to a depth of 2.5 mm over a span of 15 minutes, followed by a stabilization period of 10 minutes. Opto-responsive cells were then screened for, using 100‒250 ms long square light pulses; continuous firing activity on any spiking channels during this stimulus indicated a responsive cell was found. If a noticeable light-evoked response was lacking, the probe’s depth was increased by ∼100 micrometers, up to a maximum depth of 4.5 mm below the brain’s surface, until a responsive cell was found. After each probe movement, an additional waiting time of 5 minutes was allotted before the next set of screening pulses.

Once a responsive spiking channel was identified, a waiting period of 20 minutes was incorporated to minimize mechanical drift during recordings. If a light-responsive cell could be stably held, SpikeSort3D (Neuralynx) was employed to isolate the waveform, and the power level of the light source was adjusted such that this cell would exhibit a response within 10 ms in about ∼50% of presentations of 1‒2 ms wide light pulses; this near-threshold stimulation calibration is an important requirement for revealing potential changes in excitability which might otherwise be obscured by a saturating or insufficient stimulation strength. Following this optimization, the recording protocol began. Saline irrigation was consistently maintained for the craniotomy, and subjects were sporadically rewarded with sucrose via a handheld spout during screening for opto-responsive cells, but not once the recording started.

The recording protocol included a Pre- and a Post-stim epoch (**Figure 3A**, green boxes), each comprising 100 light pulses (1‒2 milliseconds wide) with 1-second intervals, primarily to assess recording stability. The central recording phase, the “Trial-stim” epoch (**Figure 3A**, blue box with **bold** outline) consisting of 1500 light pulses (1‒2 milliseconds wide) with varying inter-stimulus intervals (0.5 to 3.5 seconds), formed the basis for this study’s results. Lastly, the “Long-stim” epoch (**Figure 3A**, orange box), consisting of 25 wider pulses (100 or 50 milliseconds), was designed to differentiate between direct stimulation effects and post-inhibitory rebound.

**Figure 3:**
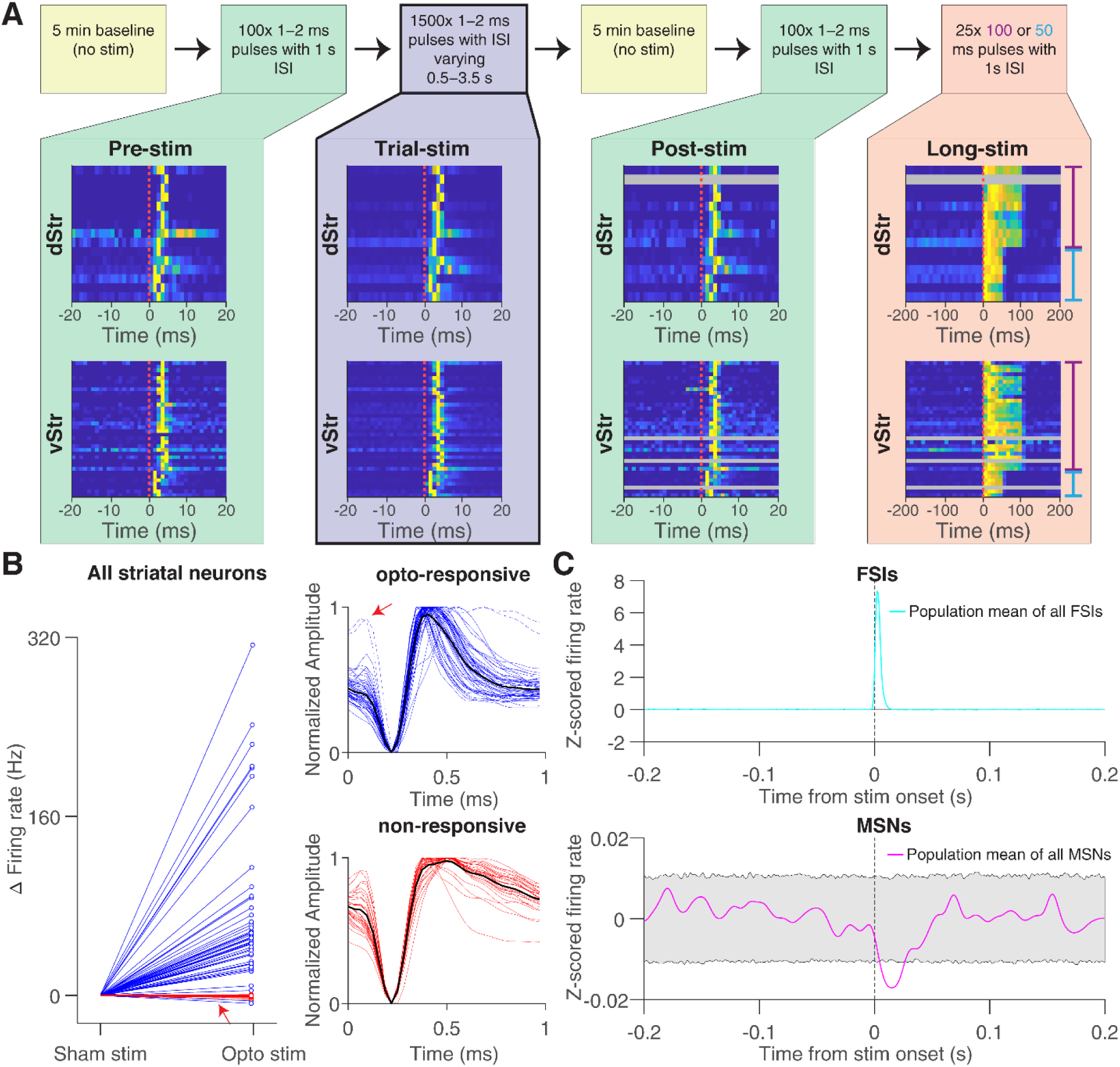
Recording protocol and characterization of ChR2-expressing cells (putative PV+ interneurons). **A**, Top: A full recording session consisted of several epochs of optical stimulation with square blue light pulses, differing in pulse widths and inter-stimulus intervals (ISI). *Pre-* and *post-stim* epochs (green) were used as baseline recordings to evaluate recording stability. The *long-stim* epoch (orange) served to distinguish between direct stimulation and post-inhibitory rebound (see main text). The *trial-stim* epoch (in blue with thick outline) constitutes the data collection epoch (1500 trials of optogenetic stimulation) that the main results are based on. Bottom: insets show averaged peri-event activity of opto-responsive PV+ interneurons, where each row represents one neuron and stimulation onset time is depicted with a dotted red line; note the clear responses to the stimulation in all neurons. In a few cases, we could not track cells through the entire recording duration at the spike sorting stage because of the mechanical drift of the optrode relative to the brain (gray rows in *post-stim* and *long-stim* epoch). These cells were included in further analysis as long as they could be reliably tracked for a minimum number of trials in the *trial stim* epoch (see *Methods: Spike-sorting* for details). **B**: Striatal cell type categorization. We categorized putative single units as either opto-responsive (blue) or non-responsive (red) based on the difference in firing rate in 10 msec windows either side of the optical stimulation (ΔFR, “post minus pre”), relative to a null distribution based on randomly chosen ‘sham’ stimulations (Kolmogorov-Smirnov 2-sample test, p < 0.01). All but two of these opto-responsive neurons displayed symmetrical biphasic spike waveforms characteristic of PV+ interneurons, whereas non-responsive neurons showed long after-hyperpolarizations typical of MSNs (Berke et al., 2004; Mallet et al., 2005). These two neurons with MSN-like waveforms showed a significant decrease in firing rate to the stimulus (dashed blue line, red arrows), likely due to synaptic inhibition from nearby PV+ interneurons. After excluding this neuron, 15 and 36 opto-responsive cells remained for analysis in dStr and vStr respectively. **C**: Population-level activity demonstrates cell-type specific responses to optogenetic stimulation. Z-scored firing rates of putative FSIs (cyan, top) show robust increases in activity following light delivery, while putative MSNs (magenta, bottom) exhibit a concurrent suppression of firing. This MSN suppression was significantly below chance levels (gray shaded region) for a brief post-stimulus period, consistent with direct inhibition from activated FSIs. Shaded area indicates +/− 2 SD of a shuffled control (see Materials and Methods).

At the end of the recording session, probes were withdrawn from the craniotomy, which was then covered with a silicon polymer (Kwik-sil or Kwik-cast, WPI), and the mouse was returned back to its home cage. Each animal underwent 2‒12 recording sessions (maximum of one per day), until the craniotomies could no longer be penetrated by the optrode. After the last recording session, mice were anesthetized using isoflurane and euthanized via carbon dioxide asphyxiation and transcardial perfusions were carried out. The brains were then extracted, fixed, and sectioned into 50-micrometer coronal slices. These sections were stained with DAPI and examined under a fluorescent microscope to verify placement of the probes.

### Spike-sorting

Spiking channels were sampled at 30 kHz and filtered within the range of 600 ‒ 6000 Hz. Probe contacts were organized into physically adjacent groups of four, each group functioning as a pseudo-tetrode. When any of the four channels within a pseudo-tetrode reached a voltage threshold, a 1 ms segment of the spike waveform from these channels was stored and given a timestamp with microsecond precision utilizing Cheetah acquisition software (Neuralynx). Spikes were automatically clustered based on their waveform properties using the KlustaKwik algorithm, followed by a manual pruning process using MClust 3.5 (A. D. Redish, University of Minnesota; Schmitzer-Torbert et al., 2005) to obtain putative single units (44 in dStr, 79 in vStr).

### Characterization of optically responsive cells

We measured the change in firing rate (ΔFR) for putative single units by subtracting the pre-stimulus firing rate in a 10 ms time window immediately preceding stimulation onset from the 10 ms time window following stimulation onset (“post minus pre”). We labeled a cell as opto-responsive if the distribution of its ΔFR obtained over all trials was significantly different from a null distribution constructed from 1000 rounds of randomly chosen “sham stimulations” (1500 times during the same epoch); and non-responsive otherwise. We used a Kolmogorov-Smirnov 2-sample test (p < 0.01) to compare the actual and null distributions, which we found to be more conservative than typical t-test and percentile based approaches.

Consistent with the expected effects of ChR2 activation, which typically leads to enhanced depolarization and increased spiking, we observed that the ΔFR for trial-stim in opto-responsive cells generally exceeded that of sham stimulation times, except for two neurons, which had significantly lower ΔFR than sham stimulation times. These neurons were putative medium spiny neurons (MSNs) inhibited by nearby PV+ interneurons, as indicated by their longer after-hyperpolarization mean waveforms compared to other opto-responsive striatal neurons (**Figure 3B**, top right; compare dashed vs. bold lines). We analyze these neurons separately from the main analysis in Figure 8 below.

When we examined the population-level activity, the z-scored firing rates revealed distinct response patterns between cell types. The population of FSIs showed a clear increase in firing rate following optical stimulation, while the MSN population exhibited a concurrent decrease in firing rate that fell below chance levels for a brief post-stimulus period (**Figure 3C**). These population plots were generated by computing the mean of z-scored peri-event time histograms (PETHs) across all cells of each type (FSIs and MSNs) for all the stimulation trials. Chance levels (shown as shaded regions) were determined through a shuffling procedure, where for each of 1000 rounds of shuffling, we calculated mean population PETHs using randomly selected sham stimulation times for each cell, then derived 95% confidence intervals from this null distribution of population responses.

PV+ interneurons are known to have narrower spike waveforms compared to MSNs (Berke et al., 2004; Mallet et al., 2005). Accordingly, we found narrower average spike-widths (defined by the time between the peak and trough) for the opto-responsive cells compared to the non-responsive striatal cells (opto-responsive spike-width: 0.19 ± 0.04 ms, non-responsive spike-width: 0.27 ± 0.05 ms). The peak response of these opto-responsive cells typically occurred 2‒6 ms after the onset of optical stimulation (**Figure 3A**, heatmaps in the blue box with bold outline), in accordance with previous work (Muñoz et al., 2014). This response delay appears to be due to the kinetics of ChR2, rather than indirect mechanisms such as post-inhibitory rebound; we verified this using longer stimulation epochs, which resulted in sustained firing consistent with direct ChR2 activation (**Figure 3A**, *long stim* epoch), as opposed to a transient response at stimulation offset that would indicate post-inhibitory rebound.

In a few cases, we were not able to track cells through the entire duration of the recording due to mechanical drift of the optrode relative to the brain (**Figure 3A**, gray rows in *post-stim* and *long-stim* epochs). Such cells were included only if they could be reliably tracked for at least 700 stimulation trials in the *trial-stim* epoch. In the end, our study included 51 opto-responsive cells in striatum (15 in dorsal striatum and 36 in ventral striatum), as shown in **Figure 3B**.

### LFP acquisition and preprocessing

The dorsal- and ventral-most contacts on the outermost shanks of the optrodes were designated as the LFP channels. Consistent with previous work (Berke 2009; Carmichael et al. 2017) striatal LFPs looked highly similar across multiple recording sites with minimal phase differences. Thus, when multiple viable LFP signals were acquired in a session, the one with least amount of line noise and the smallest opto-artifact was chosen for analysis. The LFP signal was either referenced to one of the reference wires/screws (preferentially to the one in the contra-lateral hemisphere) during acquisition, or both the LFP and the reference signal were acquired with reference to the ground and a post-hoc referencing was implemented by subtracting the reference-ground signal from the LFP-ground signal. LFP signals shown are all “positive-up”, i.e. peaks indicate positive extracellular potentials and valleys indicate negative potentials.

In our initial experiments, we sampled the LFP at 30 kHz, applying a band-pass filter ranging from 1 to 400 Hz. However, we noticed the frequent occurrence of an opto-electrical artifact in the LFP channels at the stimulation onset times, which extended significantly beyond the actual light stimulus (**Figure 4-1A**). We traced this artifact to the causal nature of the real-time DSP filter used during acquisition. To address this, we switched to recording the LFP with ‘Filter OFF’ settings, reducing the spread of the opto-artifact compared to the initial settings (**Extended Figure 4-1B**). Despite this adjustment, a stimulation artifact still extended beyond the duration of the stimulus, potentially interfering with LFP phase estimation at stimulus time. Therefore, we used a “causal” phase estimation method, which deliberately excluded LFP data following stimulation onset (elaborated in the *Materials and Methods: Stimulus phase estimation*). This solution enabled consistent analysis across all sessions using the same pipeline, irrespective of variations in acquisition filter settings.

**Figure 4:**
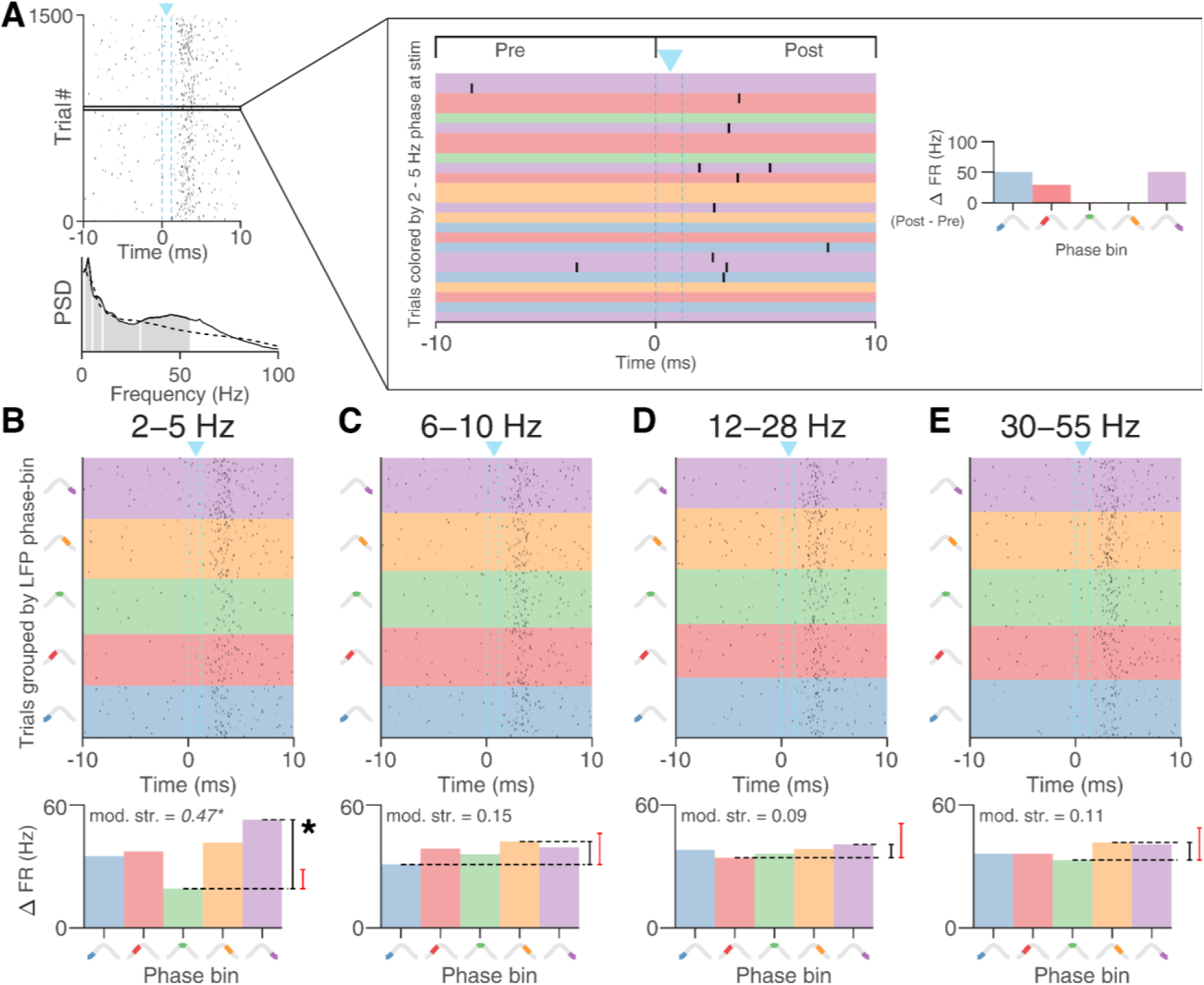
Example ventral striatal PV+ neuron with phase-dependent excitability in the delta LFP band (2‒5 Hz). **A**: Rasterplot of spiking activity around optical stimulation (dashed lines: stim onset and offset), showing a clear stimulus-evoked response. Inset: 25 trials from this session, with each row colored according to its LFP phase bin in the 2‒5 Hz band (left). Observed changes in firing rate (ΔFR, “post minus pre”) for these 25 trials are averaged across trials for each phase bin (histogram on right). This is done for all the trials of a neuron, separately for each frequency range. Bottom: Power spectrum of the local field potential (LFP) recorded simultaneously with this cell. Peaks present above the 1/f fit (dashed black line) in the delta, theta and gamma bands (indicated in gray) indicate the presence of these oscillations in the LFP signal. **B**, Top left: all trials, grouped by LFP phase bin in the 2‒5 Hz band at the time of stimulation. Evoked activity is lowest in the third phase bin (green) and highest in the first phase bin (blue) as reflected in the ΔFR vs phase-bin plot in B, bottom. The black bar indicates the modulation strength, i.e. the normalized difference between the most and least excitable phase bins (see *Materials and Methods: Modulation strength*), and the red bar indicates the chance-level difference derived from a shuffled phase distribution. The observed difference exceeds the chance difference, demonstrating significant phase-dependent excitability in the 2‒5 Hz frequency range, but not in the case of 6‒10 Hz (**C**), 12‒28 Hz (**D**), or 30‒55 Hz (**E**). LFP phases were computed using a causal filtering method to eliminate phase distortion due to the stimulation artifact (end-corrected Hilbert transform, see Extended Figures 4-1 and 4-2).

### Stimulus phase estimation

The standard method for estimating LFP phases in a given frequency band such as delta (2‒5 Hz) or theta (6‒10 Hz) is to first apply an appropriate bandpass filter for that frequency band (e.g. `filtfilt()` in MATLAB, which runs a given filter forwards and backwards to avoid phase distortion, i.e. an “acausal” filter), and then Hilbert-transforming the filtered signal to obtain the phases. However, when we employed this method to estimate the LFP phases at stimulus onset times, the distribution of these phases was markedly skewed (**Extended Figure 4-2**) when an uniform distribution is expected, indicating a problem with phase estimation. The skew in the phase distribution was not observed at time points sufficiently away from the stimulation onsets, suggesting that it results from the interaction of the acausal filtering with the stimulation artifact. One tempting solution might be to apply the Hilbert transform on only that part of the LFP signal which terminates at stimulation onset, to avoid interaction with the temporally adjacent opto-artifact. However, this approach fails because of a well-studied distortion in phase estimation at points of signal discontinuities (start or end) called Gibbs’ phenomenon (Gibbs, 1898). A solution to this problem was proposed by Schreglmann et al., 2021, called endpoint corrected Hilbert Transform (ecHT), which employs an innovative use of causal (forward) bandpass filtering for each frequency band of interest to mitigate the distortion due to Gibbs’ phenomenon.

Thus, we first divided the LFP signal into segments demarcated by consecutive trial-stim onsets, and applied ecHT separately for each frequency band of interest on each of these segments to determine the LFP phase at each sample in it. The LFP phase for a given trial was the phase of the last sample of the signal segment that terminated at the onset time of that stimulus. The use of ecHT in this way corrected the skewness of the phase-distribution at the samples corresponding to trial-stim onset, successfully eliminating the confounding influence of the stimulation artifact on phase estimation **(Extended Figure 4-2)**.

### Modulation strength

The primary dependent variable in our study, “modulation strength,” measured to what extent the response of an opto-responsive cell varied with the LFP phase during optical stimulation. Essentially, the greater the disparity in response magnitude between the most excitable and least excitable phases of stimulation, the higher the modulation strength. As before, we quantified response magnitude for a single stimulus presentation (trial) by using the difference in firing rate (ΔFR) within 10 millisecond windows before and after the onset of stimulation (“post minus pre”). A neuron exhibiting higher excitability at the time of optical stimulation would consequently show a larger ΔFR.

To determine how ΔFR depended on the LFP phase, we assigned each stimulus presentation (i.e. a single trial) a LFP phase, as calculated by the end-corrected Hilbert transform (ecHT) as described above. We then grouped all trials into five uniformly spaced phase-bins spanning −π to π radians, and calculated the mean ΔFR for each phase bin. After identifying the phase bins with the highest and lowest ΔFR as *max-bin* and *min-bin* respectively, we defined modulation strength as:

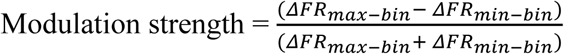

In other words, modulation strength is a contrast (difference divided by the sum) of excitability between the least excitable and most excitable LFP phases. We recognized an opto-responsive cell as having **significant phase-dependent excitability** in a specific frequency band if the z-score of its modulation strength exceeded 2 (corresponding to p = 0.022) relative to a null distribution of 1000 resampled modulation strengths. A single resampled modulation strength was obtained by randomly permuting the original phase assignments, and then calculating the modulation strength for the resulting resampled data.

### Phase locking

Beyond describing the relationship between LFP phase and response to optical stimulation, we also sought to quantify endogenous spike-LFP phase locking in opto-responsive cells. We employed the phase-locking value (PLV; Lachaux et al., 1999), mathematically represented as

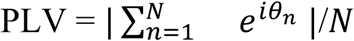

where *N* is the total spike count and *θ_n_* is the LFP phase corresponding to the sample most temporally proximal to the *n*^th^ spike. We used the phases that were obtained using the end-corrected Hilbert Transform (ecHT) method described in section *Materials and Methods: Stimulus phase estimation* to ensure uniformity between the phase-locking and the phase-dependent excitability analyses. We considered the spikes in the “trial-stim” epoch, excluding spikes that occurred 250 ms post-optical stimulation to minimize the influence of the stimulation on endogenous spike-phase locking. Only neurons with over 200 spikes in this epoch were included, resulting in 43 opto-responsive cells for analysis. To account for PLV’s known spike-count bias (Vinck et al., 2012; Aydore et al., 2013), we adopted the following subsampling procedure:

i. Randomly select 200 spikes and calculate the PLV for this subset.
ii. Repeat 1000 times, with the final PLV estimated as the mean of these 1000 values.

A neuron was then considered significantly phase-locked in a specific frequency band if the z-score of its PLV exceeded 2 (p-value < 0.022), relative to a null distribution where the spike-times were circularly shifted 1000 times with respect to the LFP signal.

## Results

We sought to determine if the excitability of striatal fast-spiking interneurons varied according to the phase of ongoing striatal LFP oscillations (see **Figure 1** for a schematic of this idea). For this purpose, we used an optical stimulation experiment in head-fixed mice that expressed ChR2 in PV+ interneurons (**Figure 2A, B**). After verifying that the striatal LFP contained clear oscillations in previously documented frequency bands (**Figure 2C**) we applied optical stimulation across various striatal LFP oscillation phases, and tested if the magnitude of the evoked response (change in firing rate) depended on the phase at which the stimulation occurred.

### LFP phase-dependent excitability in striatal PV+ interneurons was observed in delta, theta, beta and gamma bands

We operationalized excitability as the change in an opto-responsive cell’s firing rate (ΔFR) in 10 ms windows before and after optical stimulation, where a greater ΔFR value indicates higher excitability. We treated each optical stimulation event as an individual trial (*trial stim* epoch, **Figure 3A**, blue box), with a corresponding ΔFR and LFP phase (as determined at stimulus onset, see *Materials and Methods: Stimulus phase estimation*). We grouped the trials into five uniformly spaced phase bins, and calculated the mean ΔFR for each bin. We devised a measure called “**modulation strength**”: the (normalized) difference between the mean ΔFR of the most excitable and least excitable phase bins, to capture the relationship between LFP phase and excitability (*Materials and Methods: Modulation strength*). We focused on oscillations in the delta (2‒5 Hz), theta (6‒10 Hz), beta (12–28 Hz) and gamma (30‒55 Hz) bands, as they are consistently present in the striatal LFP, as confirmed by power spectral density plots (**Figures 4A** and **2C**). Thus, we proceeded to investigate whether the excitability of the striatal PV+ interneurons depended on the phases of these LFP oscillations.

We categorized a cell as **significantly phase-dependent excitable** in a specific frequency band if the z-score of its modulation strength in that band, relative to a null distribution, exceeded 2. **Figure 4** shows one such cell recorded in vStr, demonstrated significant phase-dependent excitability in the 2‒5 Hz band (**Figure 4B**; note that the maximum difference in mean ΔFR among the phase bins, blue minus green, exceeds the chance levels determined by the shuffle procedure), but not in other frequency bands. **Figure 5** shows a few other example cells that show significant phase-dependent excitability in the other frequency bands: 6‒10 Hz (**Figure 5A**), 30‒55 Hz (**Figure 5C**) and another cell that shows significant phase-dependent excitability in multiple bands (**Figure 5B**; 2‒5 Hz, 12–28 Hz, 30‒55 Hz).

**Figure 5:**
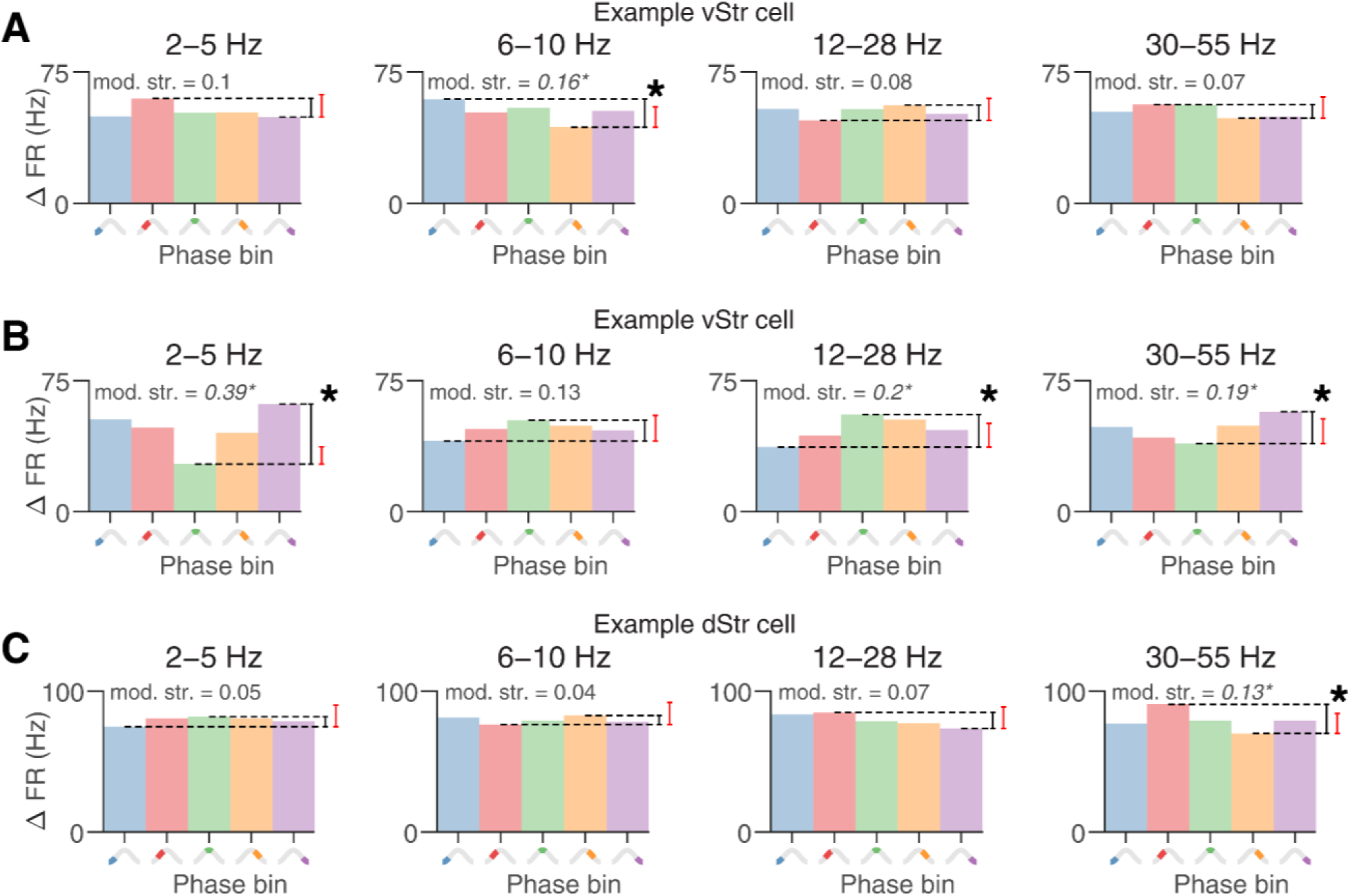
Further example ventral striatal PV+ cells with phase-dependent excitability in different frequency bands. **A**: vStr neuron with significant phase-dependent excitability in the 6‒10 Hz band. **B**: A different vStr neuron with significant phase-dependent excitability in the 2‒5 Hz, 12–28 Hz and 30‒55 Hz bands. **C**: A dStr neuron with significant phase-dependent excitability in the 30‒55 Hz band. As in Figure 4, black bars indicate modulation strength, i.e. the normalized difference between the most and least excitable phase bins, and the red bar indicates the chance-level difference derived from a shuffled phase distribution. Asterisks denote significant phase-dependent excitability.

We next characterized phase-dependent excitability at the population level. Considering the relatively small number of opto-responsive cells (51) and their unbalanced distribution between dStr (15) and vStr (36), we examined them collectively as striatal interneurons, without distinguishing between dorsal and ventral subdivisions. We noted a near significant correlation (R^2^ = 0.27, p = 0.05) between recording depth and z-scored modulation strength in the 2–5 Hz band, such that delta-band phase-dependent excitability was more likely in ventral recording locations; no significant correlations were found between recording locations and modulation strengths in any other frequency bands (**Figure 6A, B**). Overall, 12/51 (23.5%) of the striatal interneurons showed significant phase-dependent excitability in the 2‒5 Hz band, 2/51 (3.9%) in the 6‒10 Hz band, 6/51 (11.8%) in the 12–28 Hz band, and 11/51 (21.6%) in the 30‒55 Hz band (**Figure 6C**). The modulation strengths was the highest in the 2‒5 Hz band (mean = 0.22 ± 0.12, n = 12), the lowest in the 6‒10 Hz band (mean = 0.08 ± 0.02, n = 2), and intermediate in the 12–28 Hz (mean = 0.14 ± 0.07, n = 6) and 30‒55 Hz bands (mean = 0.12 ± 0.1, n = 11). Additionally, 6/51 neurons exhibited phase-dependent excitability in multiple frequency bands simultaneously. In sum, over a third of the sampled interneurons (21/51, 41.2%) showed significant phase-dependent excitability in at least one of the frequency bands.

**Figure 6:**
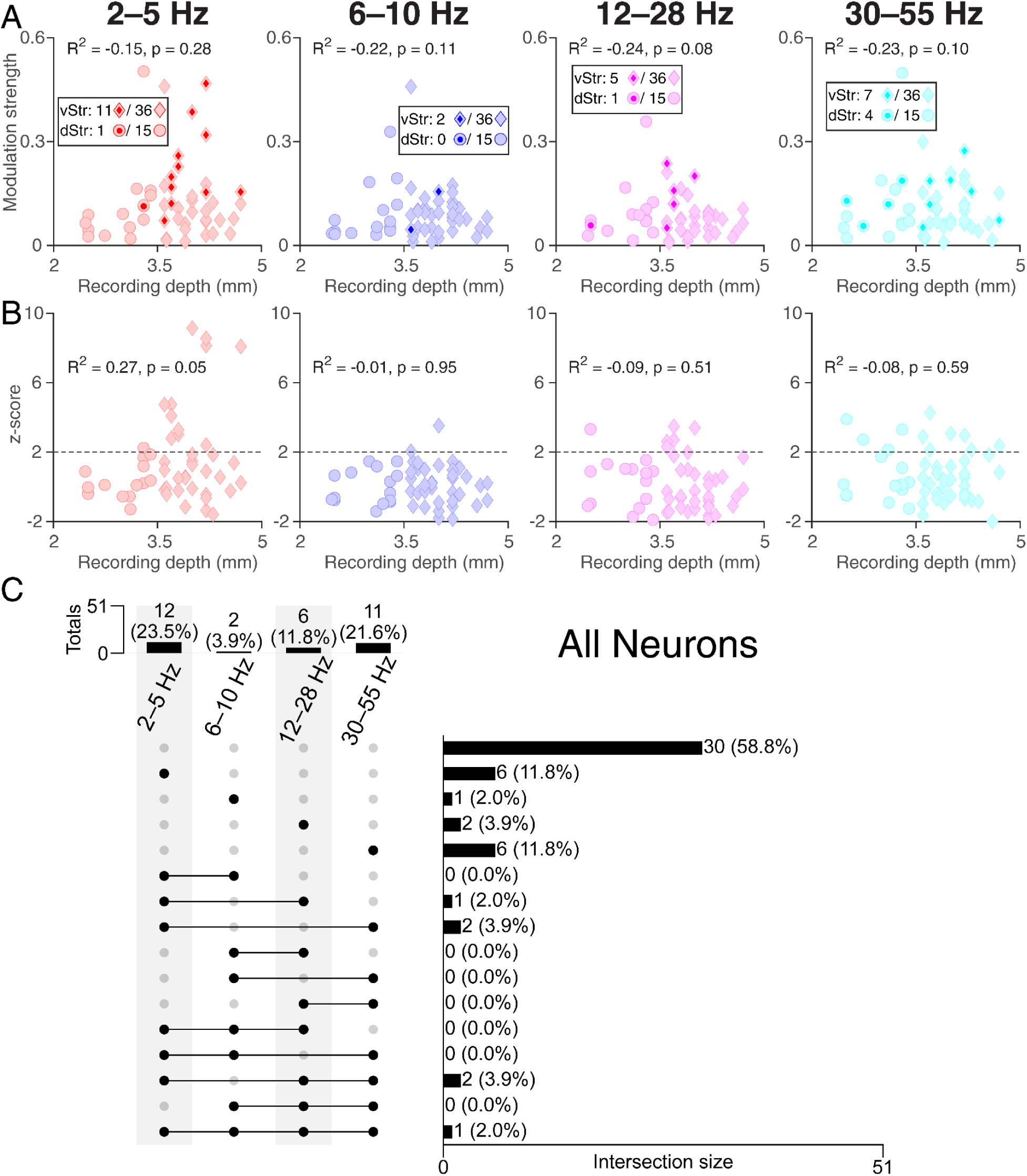
Phase-dependent modulation of all cells across frequency bands and recording depths. **A**: Scatterplots illustrate the relationship between recording depth and modulation strength for all opto-responsive cells. Circles represent dorsal striatal (dStr) neurons, while diamonds indicate ventral striatal (vStr) neurons. Darker, smaller shapes inside the larger, lighter shape signify neurons with significant phase-dependent excitability. In the 2‒5 Hz band (left), 12/51 (23.5%) opto-responsive neurons showed significant phase-dependent excitability, 2/51 (3.9%) in the 6‒10 Hz band (middle-left), 6/51 (11.8%) in the 12–28 Hz band (middle-right) and 11/51 (21.6%) in the 30‒55 Hz band (right). **B**: Scatterplot comparing recording depths with z-scored modulation strengths. A z-score above 2 (dashed line) indicates significant phase-dependent excitability in that frequency band. A borderline significant correlation (R^2^ = 0.27, p = 0.05) was observed between recording depths and z-scored modulation strength in the 2‒5 Hz band, but no other notable correlation was found between modulation strengths and recording depths in other frequency bands. **C**: Summary of the number of neurons across the entire striatum that were significant in each frequency range (top left) using “UpSet” visualization software (Lex et al., 2014). The lower right section details the various intersections. The presence or absence of a specific group in each row is denoted by a black or gray circle, respectively. The first row with four gray circles represents neurons that did not exhibit significant phase-dependent excitability in any of the frequency bands, the second row signifies neurons that showed significant phase-dependent excitability in the 2‒5 Hz band but not in others, and so on.

A potential issue when characterizing phase-dependent excitability is that firing rate is expected to change depending on phase even in the absence of any (optical) stimulation due to endogenous phase locking, i.e. cells firing preferentially at one phase rather than another. Because striatal interneurons show strong phase-locking (van der Meer & Redish, 2009; Berke, 2011; Howe et al., 2011), this introduces a potential confound that needs to be controlled for. Specifically, as described above, when our opto stimulus arrives, we define our dependent variable as ΔFR in post-stim window and pre-stim window difference. However, between these pre-stim and post-stim windows, the LFP phase has advanced by some (small) amount. If a given cell exhibits phase locking, i.e. endogenous spiking which depends on LFP phase, a difference between post- and pre-stimulus firing rate would be expected even in the absence of any effects of the stimulation (ΔFR_INTRINSIC_, see **Figure 7 A,B** for a schematic illustration of this issue). We next describe the procedure to control for this confound.

**Figure 7:**
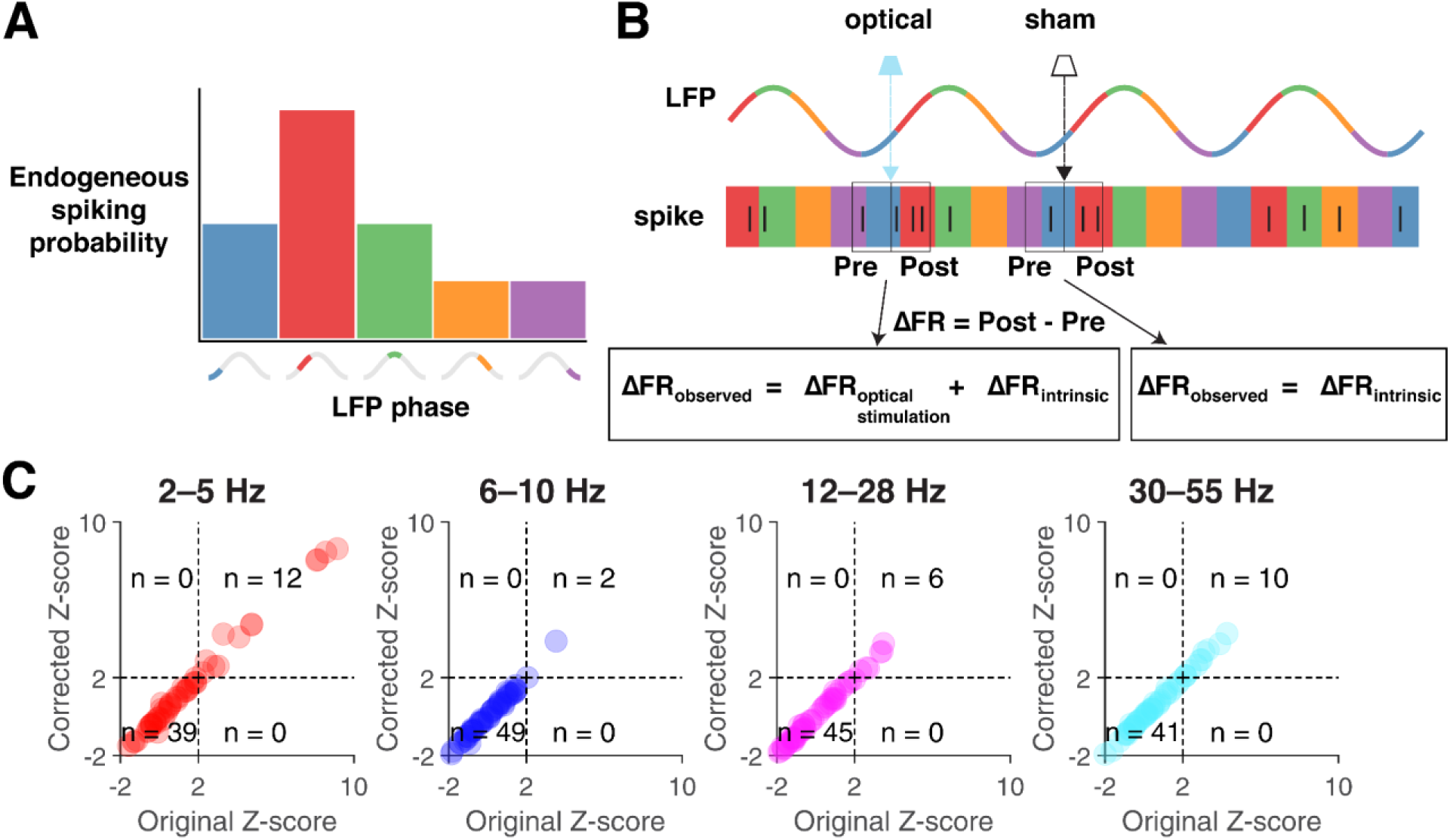
Phase-dependent excitability measure is robust to potential contribution from intrinsic spiking-phase preference. Spike-phase locking in neurons refers to a consistently higher occurrence of spiking in certain LFP phases relative to others. This results in systematic changes in firing rate as the LFP phase advances from one to the next, introducing a potential confound when estimating phase-dependent excitability. Schematically, given spiking probability as a function of the LFP phase for a hypothetical neuron (**A**) we would assess the dependence of its excitability on the LFP phase by measuring the change of firing rate (ΔFR) in 10 ms windows before (**Pre**) and after (**Post**) the occurrence of an optical stimulation (**B**). The spiking that occurs in the “**Post**” window is not just caused by the depolarization resulting from the optical stimulation, but also due to changes in spiking probability expected from the LFP phase changing (advancing to the next one) within the 10 millisecond period. Thus, observed changes in firing rate as a result of optical stimulation are potentially confounded by changes in firing rate expected from phase advancement alone. We control for this issue using the analysis described in the *Results*. **C**: Comparison of phase-dependent excitability before and after controlling for intrinsic spike-phase preference. Scatterplots show z-scored modulation strengths before (x-axis) versus after (y-axis) applying the correction. Quadrant analysis: neurons in top-right and bottom-left maintain their classification as significantly phase-dependent excitable or non-excitable, respectively. Bottom-right shows neurons that lose significance after correction, while top-left shows neurons that acquire significance. The preservation of neuronal classification across all cells demonstrates that phase-dependent excitability is robust and independent of intrinsic spike-phase preference.

We first estimated ΔFR_INTRINSIC_ for each opto-responsive neuron as a function of phase. This was done by obtaining the firing rate as a function of phase based on spikes used for quantifying intrinsic phase locking (*Materials and Methods: Phase Locking*), and then taking its derivative with respect to phase. We then applied a correction for each trial based on the phase at the time of stimulation, by subtracting the corresponding ΔFR_INTRINSIC_ from the observed ΔFR for that given trial. We recalculated and tested the modulation strength for significance (as described in *Materials and Methods: Modulation strength*) but now based on these “corrected” ΔFR values. We found that none of the cells changed its classification as phase-dependent excitable after applying this correction (**Figure 7C**). Thus, we concluded that phase-dependent excitability, as reflected by modulation strength, is not confounded by firing rate changes expected due to endogenous spiking.

While our analyses thus far have focused on ChR2-expressing PV+ FSIs, we also examined potential phase-dependent effects in medium spiny neurons (MSNs), which constitute 95% of striatal neurons and receive inhibitory inputs from local FSIs. Because MSNs tend to have low spontaneous firing rates, it is challenging to identify significant decreases in firing rate against an already low baseline. Nevertheless, among our recorded neurons, we identified two putative MSNs that met the statistical threshold for stimulus-evoked firing rate changes. These two neurons exhibited distinct characteristics from our optogenetically-identified PV+ FSIs: their waveforms lacked the characteristic biphasic profile seen in FSIs (**Figure 3B**, top right, red arrow, **Figure 8A**), and importantly, it showed a decrease rather than increase in firing rate following optical stimulation (**Figure 3B** bottom-right, **Figure 8B**). The inhibitory response in these MSNs displayed significant phase-dependence across all analyzed frequency bands, with the strongest modulation observed in the 2–5 Hz band (**Figure 8C, D**). This observation indicates that striatal MSNs can exhibit phase-dependent inhibition, presumably through phase-dependent activation of local FSIs.

**Figure 8:**
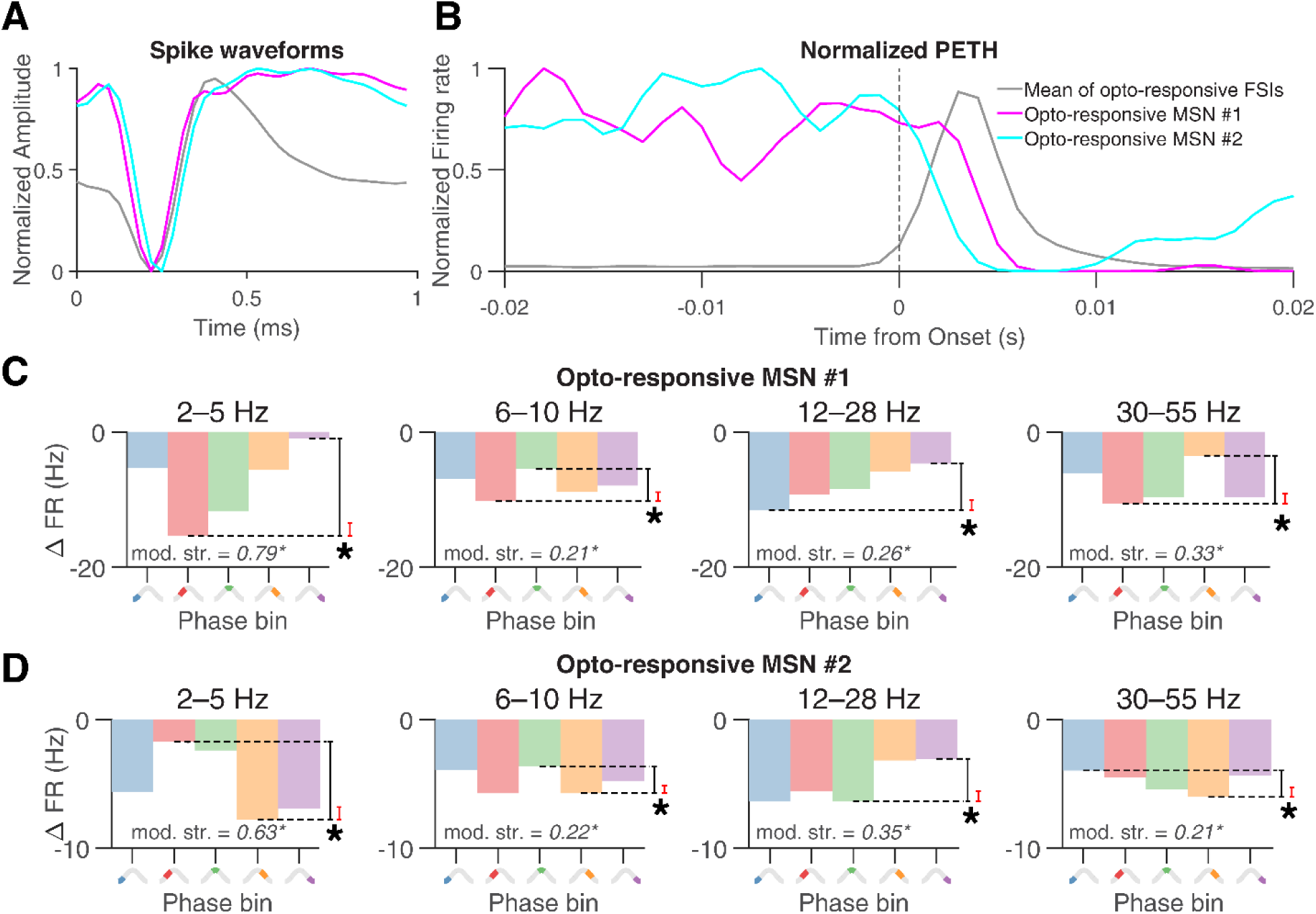
Phase-dependent inhibition in putative medium spiny neurons (MSN). **A:** Comparison of spike waveforms between two putative MSN that met the threshold for a firing rate change to stimulation (cyan and magenta; note the long afterhyperpolarization) and optogenetically identified PV+ fast-spiking interneurons (FSIs, gray). The MSNs lack the characteristic biphasic waveform seen in the mean FSI waveform. **B:** PV+ FSIs show a clear, stereotyped burst of spiking in response to optogenetic stimulation (cyan, peaking around ∼3 ms following stimulation onset). In contrast, MSN firing rate abruptly dropped following stimulation, with inhibition outlasting the peak of FSI activation (∼5‒10 ms) consistent with inhibition by FSIs. **C, D:** For these two MSNs, the magnitude of the stimulation-evoked inhibition was strongly phase-dependent, most clearly in the 2‒5 Hz frequency hand (left) but also significant in all other frequency bands. Analysis and statistics follow the same procedure as Figure 4B and Figure 5, except that ΔFR is now negative, indicating inhibition.

### Phase-dependent excitability and endogeneous spike-phase locking tend to co-occur, but are distinct phenomena

Having established the existence of phase-dependent excitability, and that it is not simply a consequence of endogenous phase-locking, we next sought to establish the relationship between these two phenomena. If the preferred excitability phase and the preferred phase-locking phase aligned, then it would be possible to infer one from the other, obviating the need for laborious experiments to determine excitability when phase-locking can be routinely measured. Thus, we first confirmed that opto-responsive cells in our study indeed exhibit endogenous spike-phase locking. We quantified phase-locking using the phase-locking value (PLV; Lachaux et al., 1999) which is equivalent to the circular mean of spikes represented by the LFP phase at which they occur, for each of the opto-responsive neurons. After excluding cells not meeting the minimum spike count for PLV (See *Materials and Methods: Phase Locking* for details), 43 cells remained. We then classified a cell as “**significantly phase-locked**” in a particular frequency band if the z-score of its PLV compared to a null distribution, exceeded 2 (p-value < 0.022). In line with prior work (van der Meer & Redish, 2009; Sharott et al., 2009), we found that a substantial fraction of these striatal PV+ interneurons were significantly phase-locked in each frequency bands: 26/43 (60.5%) in 2‒5 Hz, 21/43 (48.8%) in 6‒10 Hz, 14/43 (32.5%) in 12–28 Hz, and 21/43 (48.9%) in 30‒55 Hz (**Figure 9A**).

**Figure 9:**
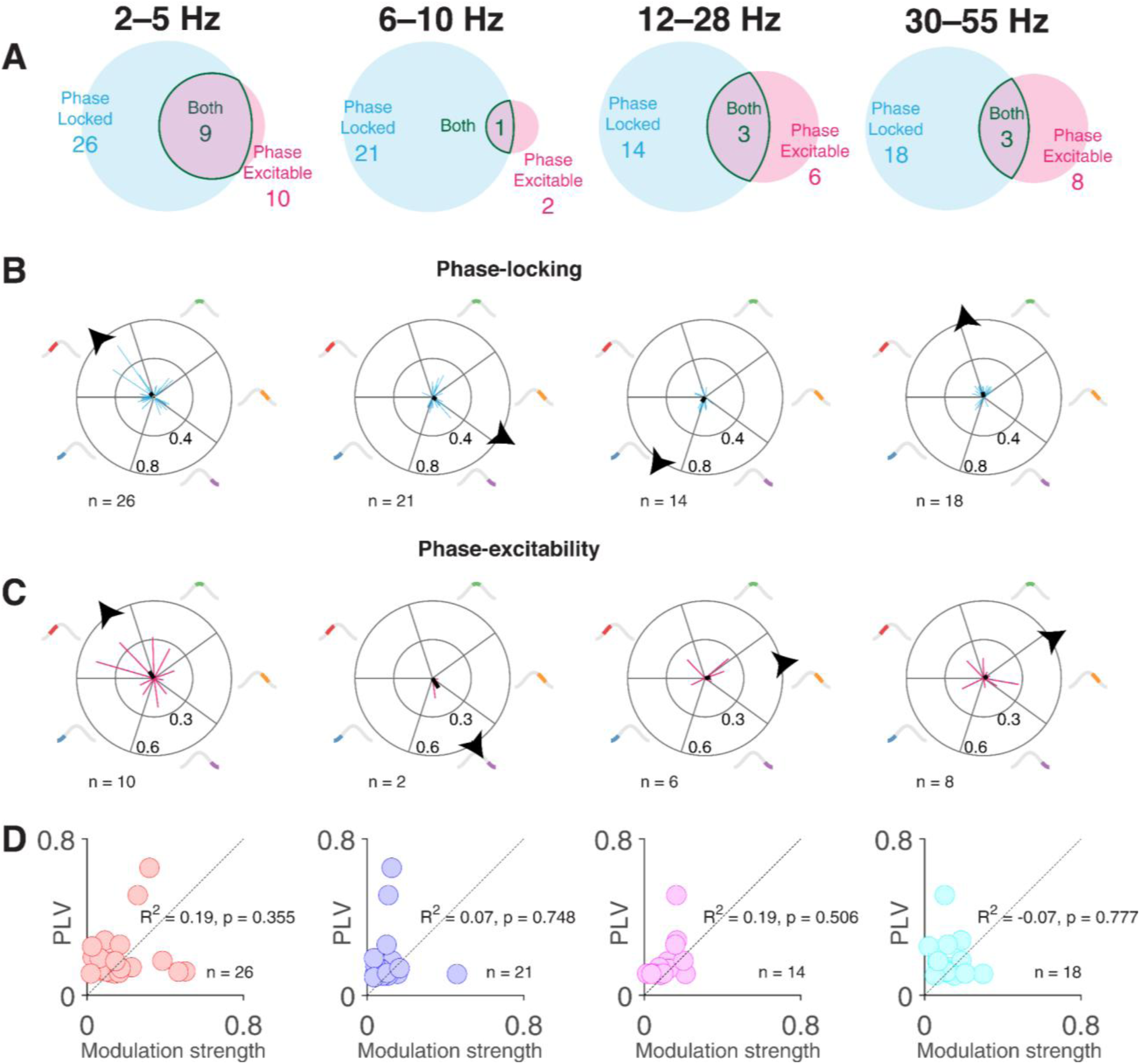
Relationship between phase-dependent excitability and endogenous spike-phase locking. Rows A through D display data across three frequency bands: delta band (2‒5 Hz) in the left column, theta band (6‒10 Hz) in the middle column, and gamma band (30‒55 Hz) in the right column. **A**: Venn diagrams illustrate the proportion of neurons with significant phase-dependent excitability (pink), significant phase-locking (gray), and their overlap (solid outline). **B**: Polar plots showcase vectors whose lengths represent the Phase Locking Value (PLV) of significantly phase-locked cells, with angles reflecting the mean phases of the LFP at spike occurrences. Individual cells are shown as blue lines, while the thick black line signifies the population’s mean vector (lengths: 0.06 for 2‒5 Hz, 0.05 for 6‒10 Hz, 0.06 for 12–28 Hz, and 0.05 for 30‒55 Hz). The *angle* of this mean vector is marked by a black arrowhead on the outer ring (2.33 radians for 2‒5 Hz, −0.52 radians for 6‒10 Hz, −2.19 for 12–28 Hz, and 1.8 radians for 30‒55 Hz). **C**: Polar plots represent vectors of neurons with significant phase-dependent excitability. Vector lengths indicate modulation strengths, and angles correspond to the centers of phase bins with the largest firing rate changes following optical stimulation. Pink lines represent individual cells, their angles slightly jittered for clarity. The population’s mean vector is depicted by a thick black line (lengths: 0.06 for 2‒5 Hz, 0.09 for 6‒ 10 Hz, 0.04 for 12–28 Hz, and 0.05 for 30‒55 Hz). The angle of this mean vector is marked by a black arrowhead on the outer ring (2.19 radians for 2‒5 Hz, −1.01 radians for 6‒10 Hz, 0.19 for 12–28 Hz, and 0.53 radians for 30‒55 Hz). The alignment between the angles of the population vectors (black arrowheads in rows B and C) in the 2‒5 Hz band was significantly less than chance (bootstrap significance test, p = 0.004), and less than chance in 30–55 Hz (bootstrap significance test, p = 0.031). but not in 6‒10 Hz (bootstrap significance test, p = 0.423), or 12‒28 Hz (bootstrap significance test, p = 0.682). **D**: No significant correlation was observed between the PLV and modulation strengths of cells that exhibited significant phase-locking and phase-dependent excitability in the individual frequency bands of interest.

Having established the prevalence of endogenous phase-locking, we next examined the relationship between significant phase-locking and phase-dependent excitability. The majority of the significantly phase-dependent excitable neurons among these 43 cells were also significantly phase-locked (**Figure 9A**): 9/10 (90%) in 2‒5 Hz, 1/2 (50%) in 6‒10 Hz, 3/6 (50%) in 12–28 Hz, and 3/8 (37.5%) in 30‒55 Hz; note that a few of the significantly phase-dependent excitable cells did not pass the minimum spike criterion needed for PLV, leading to a lower number of cells showing phase-dependent excitability than reported in the previous section. These high percentages indicate that a striatal interneuron that is significantly phase-dependent excitable in a frequency band, is also likely to be significantly phase-locked in the same band.

Given that significant phase-locking and significant phase-dependent excitability have a high degree of overlap, we then asked if the degree to which they phase-lock (PLV) is related to the modulation strength of their phase-dependent excitability. We did not find a significant correlation between the PLV and modulation strength among the significantly phase-locked neurons (**Figure 9D**) in the 2‒5 Hz (R^2^ = 0.19, p = 0.355), 6‒10 Hz (R^2^ = 0.07, p = 0.748), 12‒28 Hz (R^2^ = 0.19, p = 0.506), or 30‒55 Hz (R^2^ = −0.07, p = 0.777) bands. This suggests that magnitude of phase-locking does not offer a direct read-out of the phase-dependent excitability in the same frequency band.

Finally, we sought to determine if the preferred *phases* of phase-dependent excitability and that of endogenous spike-phase locking aligned. At a population level, the mean phases of significantly phase-locked neurons and the mean phase-dependent excitable phases were closely aligned in the 2–5 Hz band (black arrowheads in **Figure 9B-C**, left: phase-locking population vector angle = 2.33 radians; phase-dependent excitability population vector angle = 2.19 radians; bootstrap significance test, p = 0.004), as well as in the 30–55 Hz (black arrowheads in **Figure 9B-C**, right: phase-locking population vector angle = 1.8 radians; phase-dependent excitability population vector angle = 0.53 radians; bootstrap significance test, p = 0.031); but not for the 6–10 Hz (black arrowheads in **Figure 9B-C**, middle-left: phase-locking population vector angle = −0.52 radians; phase-dependent excitability population vector angle = −1.01 radians; bootstrap significance test, p = 0.423), and the 12–28 Hz bands (black arrowheads in **Figure 9B-C**, middle-right: phase-locking population vector angle = −2.19 radians, phase-dependent excitability population vector angle = 0.19 radians; bootstrap significance test, p = 0.682). For these analyses, we considered the LFP phase at the time of optical stimulation for phase-dependent excitability calculations. However, the 2–5 ms activation delay inherent to optogenetic stimulation, while negligible for low-frequency oscillations, could introduce substantial phase delays at higher frequencies (e.g., 0.38 to 0.95 radians at 30 Hz). We therefore applied a frequency band-specific correction to the excitable phase calculations. After this correction, the general pattern of phase alignment remained consistent: significant alignment persisted in both the delta (2–5 Hz; phase-locking = 2.33 radians, corrected excitability = 2.27 radians, p = 0.003) and gamma bands (30–55 Hz; phase-locking = 1.80 radians, corrected excitability = 1.46 radians, p < 0.001), while remaining non-significant in theta (6–10 Hz; phase-locking = −0.52 radians, corrected excitability = −0.83 radians, p = 0.259) and beta (12–28 Hz; phase-locking = −2.19 radians, corrected excitability = 0.63 radians, p = 0.912) bands. Taken together, these results suggest that the preferred phases of endogenous spike-phase locking in delta (2–5 Hz) and gamma (30–55 Hz) frequency bands are also the most excitable.

## Discussion

In this work, we tested whether the excitability of striatal PV+ interneurons is dependent on the phase of striatal LFP oscillations. The existence of such phase-dependent excitability is a key requirement for striatum to implement flexible communication via oscillatory synchrony, as envisioned in theories such as communication through coherence (CTC). CTC proposes that an upstream region can increase its effectiveness in driving its target by aligning the timing of its inputs with the maximally excitable phases of the target. We found that approximately a third (41.2%) of striatal neurons sampled demonstrated significant phase-dependent excitability in at least one LFP frequency band (23.5% in delta, 3.9% in theta, 11.8% in beta and 21.6% in gamma); the rest (58.8%) did not meet our thresholds for significant phase-dependent excitability. Thus, this study demonstrates for the first time that a key physiological variable in the striatum – excitability, operationalized as the response to a fixed stimulus – varies rhythmically with the phase of the striatal field potential.

Importantly, we rule out two major sources of possible confounds. First, we needed to ensure that the phase estimate at the time of stimulation was not affected by stimulation artifacts. This technical issue required using a so-called “causal” filter, which unlike commonly used “acausal” filters (such as MATLAB’s `filtfilt()`) do not use data points in the future to construct the filtered signal at the current time. The more challenging step to solve was how to prevent distortions in the phase estimate at the time of the stimulation, i.e. when no future time points are available to inform the phase estimate. To do this, we used the end-corrected Hilbert Transform (ecHT; Schreglmann et al., 2021) to avoid phase distortions, as verified by the flat phase distributions that resulted (**Extended Figure 4-2**). The second possible confound is the phase-dependent change in response to our stimulus that would be expected from phase locking alone (**Figure 7**); once the logic of this concern is clear, it is relatively straightforward to compute a corrected measure which did not change our results.

The present study addresses a gap in the literature by establishing the existence of phase-dependent changes in excitability in the striatum, a fundamental requirement for flexible information routing or gating schemes such as communication-through-coherence (CTC). This finding lends at least some plausibility to the many studies that have inferred communication with striatum based on LFP coherence (DeCoteau et al., 2007; Popescu et al., 2009; Gruber et al., 2009; Hunt et al., 2009; van der Meer & Redish, 2011; Cohen et al., 2012; Lemaire et al., 2012; Dejean et al., 2013; Lansink et al., 2016; Catanese et al., 2016; Amadei et al., 2017). It also is a proof of principle that a fundamental requirement for CTC to occur in the striatum, i.e. phase-dependent excitability, is satisfied; although, as we discuss below and elsewhere, even coherence and phase-dependent excitability are not sufficient for the full CTC mechanism to work. Thus, our results address a question of fundamental theoretical importance for models of striatal function: how can this downstream area of convergence dynamically select between its multiple inputs? Several influential studies have probed this question at the intracellular level (O’Donnell & Grace, 1995; Goto & Grace, 2005; MacAskill et al., 2012), but it has been challenging to relate those results to the LFP signals that are much more conveniently accessed in typical experiments as well as in the clinic.

The most closely related work to the present study is Zhang & Frohlich (2022) who used a conceptually similar approach to demonstrate phase-dependent excitability based on alpha LFP oscillations recorded from ferret post-parietal cortex. They found that pyramidal cells, but not putative PV+ fast-spiking interneurons, showed phase-dependent excitability; this difference with our study, in which PV+ FSIs did show phase-dependent excitability, could be because of the differences in microcircuitry between cortex and striatum (e.g. GABAergic collaterals from striatal MSNs to FSIs absent in cortex; Tepper et al., 2004; Berke, 2011). Surprisingly, the most excitable phase of the neurons in Zhang & Frohlich (2022) did not systematically align with their preferred phase-locking phase. In contrast, in our study, there was clear alignment between phase-locking and excitability at least in the delta and gamma bands, although even in that case, not every individual neuron had aligned phases, and the magnitude of endogenous phase-locking did not correlate with the magnitude of phase-dependent excitability The relationship between phase-locking and phase-dependent excitability likely depends on multiple factors, and may do so in nonlinear ways: for instance, if a neuron is driven hard by excitatory input at a specific LFP phase, then additional input may be ineffective due to low input resistance, adaptation, and/or refractoriness. Conversely, if a neuron is strongly inhibited (e.g. in a down-state) then inputs may also be ineffective, but for a very different reason. More generally, phase-locking may be affected by changes in afferent activity (e.g. firing rate or timing changes in feedforward excitatory input), and/or by changes in local excitability (e.g. a “gating” mechanism such as local inhibition, shunting, up/down states, et cetera). Thus, we should not necessarily expect a direct correspondence between these two phenomena, but rather seek to characterize their relationship under different conditions. Our results offer a helpful simplification, that in the delta and gamma bands, at least the preferred phases for phase locking and excitability align.

There are several important limitations to this study. First, it is not immediately clear whether the prevalence (∼one-third of PV+ neurons) and strength (depth of modulation across phases) of phase-dependent excitability is in fact sufficient to support a scheme like CTC. In other words, how much effective gain control or “switching” between inputs would the observed amount of phase-dependent excitability support? Computational modeling studies, such as those by Akam & Kullmann (2010, 2012) go some way towards addressing this question. Although they did not simulate striatal neurons and microcircuits specifically, their conclusion that large amounts of phase-dependent modulation are required to make CTC effective sounds a cautionary note. A second limitation is that while phase-dependent excitability is absolutely necessary for CTC-like schemes to work, it is not sufficient. Most prominently, unless there is some kind of switching mechanism that can shift the phase or frequency of vStr oscillations, “communication through coherence” might equally well be described as “coherence through communication” (Schneider et al., 2021). Finally, since our focus was on establishing the basic phenomenon of phase-dependent excitability, our study was not powered to examine potential differences between striatal subregions such as dorsal vs. ventral striatum.

Our study focused on FSIs for several reasons. First, compared to MSNs, these neurons show stronger spike-field phase locking (van der Meer et al. 2010; van der Meer et al. 2019). Crucially, although FSIs are only a minority of striatal neurons (2–5%; Tepper et al. 2018) they are the mechanistic underpinning of feedforward inhibition in the striatum, exerting a powerful influence over a surprisingly large population of MSNs, as indicated by anatomical and functional studies (Gittis et al. 2010; Tepper et al. 2018). For instance, FSI activity modulates the expression of habitual behaviors (O’Hare et al. 2017) and manipulation of FSIs is sufficient to prevent compulsive grooming (Mondragon-Gonzalez et al. 2024). Thus, FSIs are in a position to serve as the “conductors” of striatal population activity patterns that underlie learned behaviors. Consistent with this idea, we find that the two MSNs in our dataset that showed significant inhibition following the optical stimulation, presumably through feedforward inhibition from nearby opto-responsive FSIs, also exhibited significant phase-dependent inhibition across all frequency ranges (Figure 8). A more general test of this idea would require a task that consistently drives MSNs to sufficiently high firing rates to be able to measure feedforward inhibition; low resting firing rates made this challenging to do in the present study. Nevertheless, the demonstrated, powerful role of FSIs in shaping striatal population activity and the proof-of-principle that MSNs can show phase-dependent excitability supports the generality of our results.

Experimentally, our proof-of-principle results suggest several productive next steps. A more physiological input stimulus, such as the stimulation of specific striatal inputs from e.g. hippocampus or prefrontal cortex, would provide a more realistic test of the extent to which inputs from different sources, and their potentially different rhythmic frequency bands, are subject to phase-dependent excitability. In parallel, it remains to be tested whether observed periods of rhythmic LFP coherence are in fact associated with changes in information transfer, e.g. is striatal phase-locking or precession to the theta rhythm the result of oscillatory coherence in the theta band? Causal manipulations, across two areas, involving imposing coherent rhythms at a preferred and non-preferred phase would be particularly powerful approaches to test the possibility of CTC in the striatum under physiological conditions (Ozawa et al., 2020). Finally, it is likely that behavioral state plays a role in modulating phase-dependent excitability. For instance, in the present experiment mice were head-fixed, not engaged in any particular task, and rarely ran on the running wheel; active running or task-engagement might be expected to change e.g. theta-band excitability changes. In any case, the present work provides a proof-of-principle foundation for such next steps.

## Acknowledgments

We thank Andrew Alvarenga for constructing and maintaining the head-fixed setup and Dwayne “Whitey” Adams for manufacturing not only the probe holder, but crucially also the probe holder holder. Earlier versions of this work were presented at the SfN Annual Meeting in 2019 (Carmichael & van der Meer, “A physiological basis for communication through coherence in the rodent striatum”, SfN Abstracts 605.17) and 2022 (Mohapatra et al., “Can the rodent ventral striatum communicate through coherence?” SfN Abstracts 064.13).

## Data and code availability

Preprocessed data files are available on: https://gin.g-node.org/manimoh/vstr_phase_stim Associated analysis code is available on: https://github.com/vandermeerlab/mm_phase_stim

## Extended Data and Figures

**Figure 4-1:**
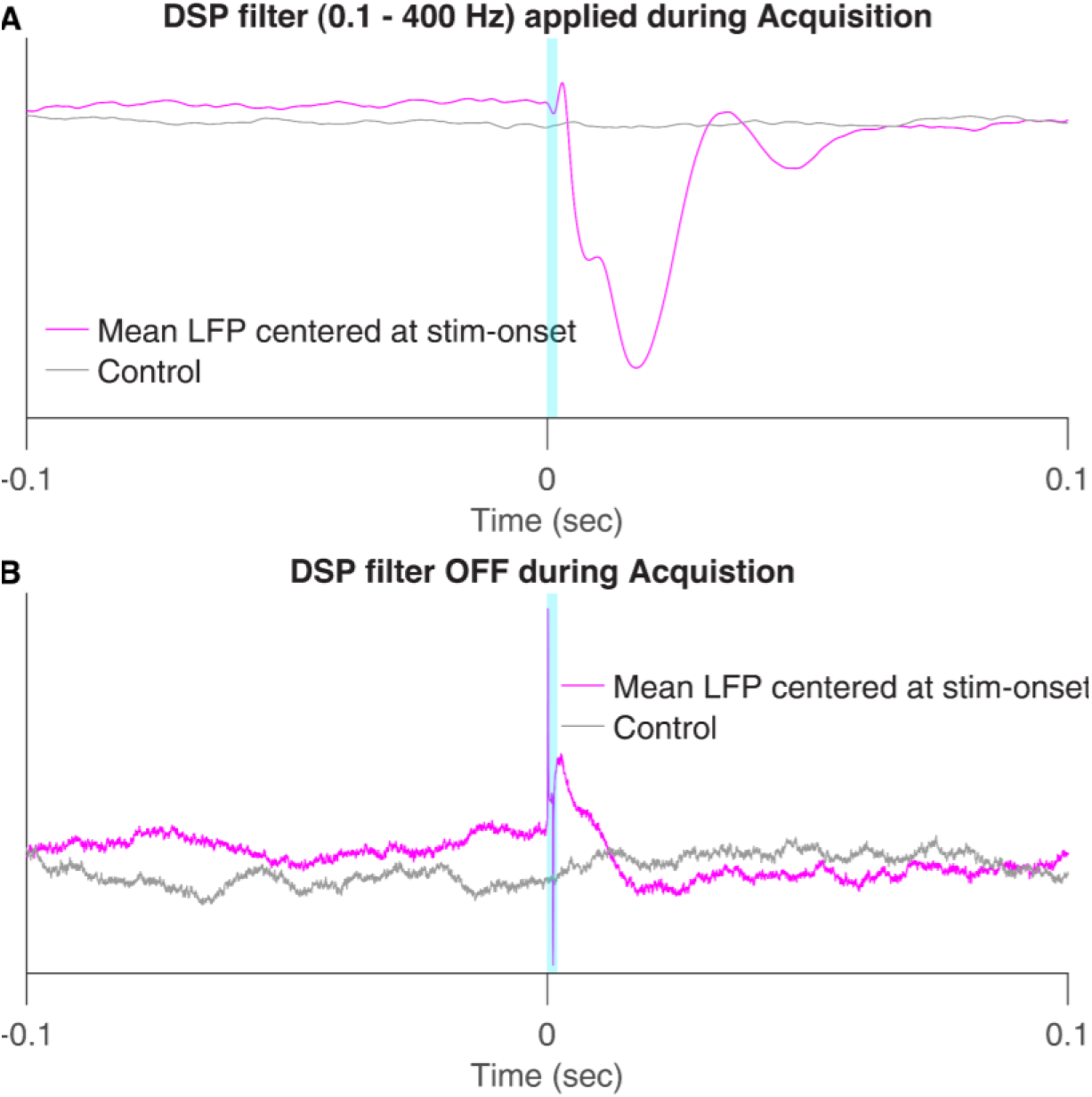
Opto-electric artifact in LFP channels. We observed a frequent occurrence of opto-electric artifacts near the stim onsets in the LFP signal in most of our recordings. A depicts one such example session where the LFP signal was recorded with a real-time DSP filter (0.1‒400 Hz) applied during acquisition. Here, we contrast the mean of all the LFP signal snippets in a 0.2 second window centered at each of the stimulus onsets, with the mean of an equal number LFP snippets which are centered around randomly chosen timepoints from the same recording epoch as the stimulation (**magenta** vs **gray**). It is noteworthy that the temporal spread of this opto-electric artifact far exceeds the pulse-width used in the light stimulation (1‒2 ms, **cyan**), which we suspected was the result of the causal nature of the real-time DSP filter. **B** is an example recording session with no DSP filters applied during acquisition where even though the LFP signal does contain opto-electric artifacts, its temporal spread is relatively less.

**Figure 4-2:**
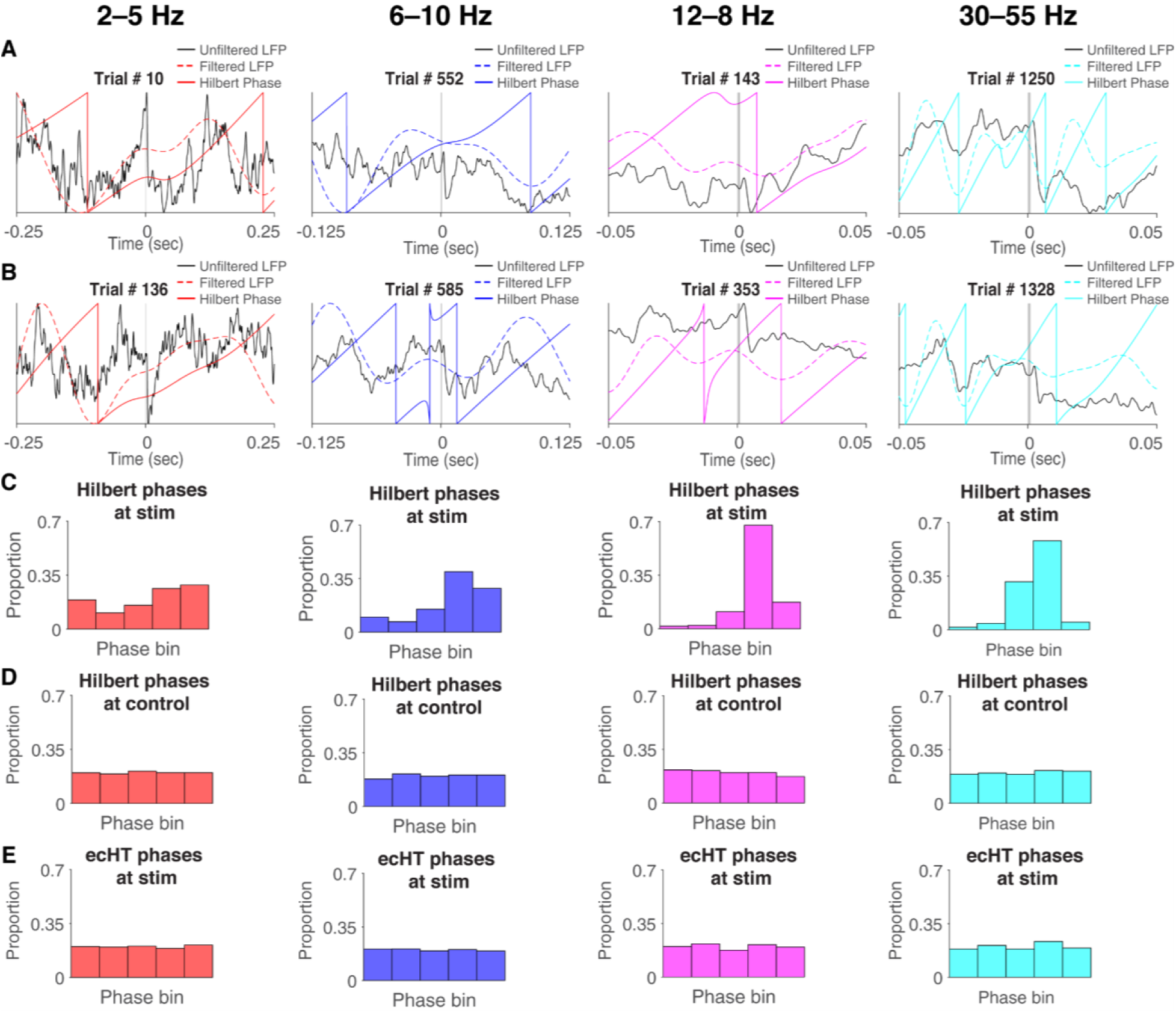
Distortion in Hilbert-phase estimation at stim-times and subsequent correction by causal phase estimation method. Rows **A** through **E** present data across four frequency bands: delta (2‒5 Hz) in the left column, theta (6‒10 Hz) in the middle-left, beta (12–28 Hz) in the middle-right, and gamma (30‒55 Hz) in the right. In neuroscience, estimating the phases of LFP oscillations at specific times typically involves applying an acausal bandpass filter to the LFP signal, followed by a Hilbert transform to extract the phase at required time points. However, during optical stimulation, light artifacts in the LFP signal can distort Hilbert phase estimation near stimulation times. Panels **A** and **B** show example trials from each frequency band in a representative session, highlighting this distortion near the stim onset. Unfiltered LFP traces at stim onset are shown in solid black. Corresponding filtered LFP traces and their resultant Hilbert phases are represented by solid and dashed colored lines (**red** for delta, **blue** for theta, and **green** for gamma). All traces are min-max normalized for clarity, with the optical stimulation period marked in cyan. Given that stim onsets were predetermined and intervals between them varied (0.5‒3.5 s), an even distribution of phases at these points was anticipated. However, applying this method to the session depicted, we observed a skewed distribution of phases at stim onset times across all bands (**C**), but not at random time points within the same epoch (**D**). This skew was attributed to interference from opto-artifacts in the LFP signal with acausal filtering and Hilbert transformation. Employing end-corrected Hilbert transform (ecHT) on signal snippets ending at stim onset effectively eliminated this skew in phase distribution (**E**). The use of ecHT, as detailed in *Materials and Methods: Stimulus phase estimation* provides a more accurate estimation of LFP phases at optical stimulation onsets, free from the distortions caused by optical stimulation artifacts.

## Footnotes

1 Following convention, we use the abbreviation LFP here to refer to the field potential, even though we acknowledge that the striatal LFP is “not so local”, i.e. contains prominent volume-conducted components from other brain structures (Carmichael et al., 2017; Lalla et al., 2017; Carmichael et al., 2019).

